# Postnatal development of skeletal muscle in IUGR pigs: morphofunctional phenotype and molecular mechanisms

**DOI:** 10.1101/524629

**Authors:** A.D. Pereira, F. Felicioni, A.L. Caldeira-Brant, D. Magnabosco, F.P. Bortolozzo, S.C. Tsoi, M. K. Dyck, W.T. Dixon, P.M. Martinelli, E.C. Jorge, H. Chiarini-Garcia, F.R.C.L. Almeida

## Abstract

Intrauterine growth restriction (IUGR) is a serious condition which impairs the achievement of the fetus full growth potential and occurs in a natural and severe manner in pigs. Knowledge on skeletal muscle morphofunctional phenotype and its molecular regulation in IUGR pigs is important to understand postnatal muscle development and may help the establishment of therapies to improve skeletal muscle growth in those individuals. To investigate the impairment of skeletal muscle postnatal development due to IUGR, we evaluated the histomorphometrical pattern of the semitendinosus muscle, the Myosin Heavy Chain (embryonic, I, IIa, IIb and IIx MyHC) fiber composition and the relative expression of genes related to myogenesis, adipogenesis and growth during three specific periods: postnatal myogenesis (newborn to 100 days of age), postnatal development (newborn to 150 days of age), and hypertrophy (100 days to 150 days of age), comparing IUGR and normal birth weight (NW) pigs. Growth restriction in utero affected muscle fiber diameter, total fiber number and muscle cross sectional area which were smaller in IUGR pigs at birth (P < 0.05). Even though the percentage of MyHC-I myofibers was higher in IUGR females at birth (P < 0.05), in older gilts, a lower percentage of MyHC-IIx isoform (P < 0.05) and the presence of emb-MyHC were also observed in that experimental group. Regarding the pattern of gene expression in the postnatal myogenesis period, growth restriction in utero led to a down regulation of myogenic factors, which delayed the expression of signals that induces skeletal muscle myogenesis (*PAX7, MYOD, MYOG, MYF5* and *DES*). Taken together, the muscle morphofunctional aspects described and their ontogenetic regulation define the possible molecular origins of the notorious damage to the postnatal musculature development in IUGR pigs.

## Introduction

Intrauterine growth restriction (IUGR) is defined as the impaired development of the mammalian fetus, or its organs, preventing it from reaching its full growth potential. This condition is characterized by low birth weight and is considered the second cause of infant mortality in the world [1]. It is also associated to the predisposition to certain chronic diseases (e.g. hypertension, obesity and diabetes) in adulthood. The main cause of IUGR is an insufficiency of the placenta in distributing enough nutrients and oxygen to the offspring [2].

IUGR is a significant problem not only in human neonatology but also in swine production. Breeding selection for increased litter size in this species has resulted in increased number of small piglets at birth, in particular those affected by IUGR [3] [4] [5]. In this species, growth restriction *in utero* is mainly caused by uterine crowding, resulting in asymmetrical IUGR [6] [7]. Moreover, this condition may increase the risk of neonatal morbidity or affect the piglets’ postnatal growth performance [8] [9].

Recently, it was shown that proteins related to energy supply, metabolism and structure, function and proliferation of the skeletal muscle cells were differentially expressed in IUGR-affected piglets [10]. For these reasons, IUGR is related to economic problems for meat production, such as reduced feed conversion efficiency, decreased percentage of meat [11] and increased percentage of body fat in the carcass [12]. Additionally, the pig provides an excellent animal model to translational medicine implications of IUGR, as it shows similarities with humans regarding anatomy, metabolism, and rapid postnatal growth rate [7] [13] [14].

There is evidence that postnatal characteristics determined by IUGR are programmed during intrauterine development [15]. Compared to brain and heart, skeletal muscle and adipose tissue have a lower priority for nutrients repartitioning, which makes these tissues especially vulnerable to nutritional deficiency *in utero* [16]. Thus, IUGR individuals exhibit compromised postnatal growth [17] [18] which may be due to skeletal muscle damage [19] and delayed skeletal muscle maturity [20].

At the molecular level, skeletal muscle development during postnatal life is dependent on prenatal myogenesis [21]. In the vertebrata clade, molecular factors involved in myogenesis participate hierarchically in both intrauterine and postnatal muscle development [22] [23]. During the prenatal period, cells recruited to the myogenic lineage are the embryonic muscle progenitors, while in the postnatal period, these progenitor cells are quiescent, located at the periphery of the muscle fiber, and referred to as satellite cells [21] [24]. These later cells are of great importance for skeletal muscle myogenesis in the postnatal period. Under the same genetic hierarchy that determines embryonic myogenesis, satellite cells are recruited to the myogenic lineage by the *PAX7* gene, giving rise to myoblasts. On the other hand, some myoblasts can eventually return to the quiescent state (satellite cells) or the expression of *MYF5* and *MYOD* commit cells to the myogenic program. Finally, the expression of *MYOG* (Myogenin), a terminal differentiation gene, will characterize the muscle differentiation state: formation of myotubes/myofibers [21] [24].

In this context, knowing that IUGR imposes a limitation on skeletal muscle postnatal development, it is imperative to characterize the morphological modifications and molecular mechanisms that govern it. This knowledge may reveal the biological targets for future therapeutic interventions to improve postnatal myogenesis in IUGR individuals.

## Materials and Methods

### Animals and experimental design

Sixty newborn female pigs DB-DanBred genotype (crossbred between Landrace and Large White breeds) from 30 litters, born to 4^th^ - 6^th^ parity sows, in litters of 10 to 15 total born piglets were selected immediately after birth (before they had suckled colostrum) and were divided into two birth weight categories: normal weight (NW: birth weight range from 1.4 to 1.7 kg; n=30) and intrauterine growth restricted (IUGR: birth weight range from 0.7 to 1.0 kg; n=30) littermates. The criteria used at selection were based on the concept of intrauterine crowding as performed in a previous study [17]. Birth weight ranges for each experimental group were determined as mean +1 standard deviation (SD) to mean +2 SD for the NW group and mean – 2 SD to mean – 1 SD for the IUGR group, based on the average (mean) and the SD of birth weights previously obtained from 1,000 newborn piglets of the same genetic line [25]. Runts, defined as piglets weighing less than 700 g, were excluded. Furthermore, in order to overcome possible litter birth weight effects on fetal development [26], the piglets selected belonged to litters whose mean birth weight ranged from 1.25 kg to 1.65 kg based on the average litter birth weight registered at the farm in the previous year.

At the end of selection, three experimental groups were obtained: one sub-set of 10 pairs of female littermates from each experimental group which was euthanized at birth (newborn – NB), one sub-set of 10 female littermates euthanized at 100 days of age (juvenile) and one sub-set of 10 pairs of female littermates from each experimental group euthanized at 150 days of age (adult). In the sub-set necropsied at birth, some organs (e.g. heart, pancreas, liver, spleen, small intestine, large intestine, kidneys, semitendinosus muscle and brain) were weighed and the occurrence of IUGR was confirmed by comparing the brain to liver weight ratio, according to Alvarenga and colleagues [17]. The other sub-sets were reared in group pens, grouped by birth weight class, until the finishing period (~150 days of age).

Feed and water were provided *ad libitum* throughout the nursery and growing-finishing phases. Pigs were fed standard nursery and growing-finishing diets expected to meet requirements for lean growth performance. Piglets were weaned at 23.1 days old on average; the nursery period lasted six weeks, and the growing-finishing phase lasted 12 weeks. Individual body weight of the females was recorded at weaning, and at the end of the nursery and growing-finishing periods, without restriction of feed and water. The experimental protocol was approved by the Ethical Committee in Animal Experimentation of the Federal University of Rio Grande do Sul (protocol #23732).

### Tissue preparation

Following euthanasia, samples of the semitendinosus muscle were taken from the muscle origin, which is the muscle extreme close to the ischiatic tuber, and were subjected to different processing steps, according to the histomorphometrical, immunofluorescence, and gene expression analysis. For histomorphometrical evaluations, samples of 1-2 mm thickness were fixed through immersion in 5% (w/v) glutaraldehyde (Biological Grade, EMS, #16500) in 0.05M phosphate buffer pH 7.3 for 24 hours, dehydrated in increasing concentrations of ethanol, embedded in glycol methacrylate plastic resin (Historesin, Leica, Heidelberg, Germany), sectioned at 3 μm thickness and stained with toluidine blue-sodium borate [27]. To perform immunofluorescence quantification of muscle fiber types, samples were fixed in 4% paraformaldehyde (w/v) in 0.05M phosphate buffer pH 7.3 for 24 hours and embedded in Paraplast (Sigma Aldrich, São Paulo, Brazil). Sections of 5 μm thickness were placed on silanized slides. For gene expression studies, fresh muscle samples were preserved in RNA holder (Bio Agency, São Paulo, Brazil) for 24 hours overnight at 4°C and subsequently stored at -20°C.

## Histomorphometrical analyses

### Muscle cross sectional area and fiber number

Before embedding, muscle samples from newborn piglets were placed on an acetate sheet and their transversal circumference drawn. These drawings were scanned and the total cross sectional area of each muscle sample calculated through an analysis program (Image J, 1.49v – free version, National Institutes of Health, Bethesda, MD, USA). Additionally, 10 randomly selected histological fields were used per animal to calculate the number of fibers per area (mm^2^ - fiber density). These values were obtained using the same analysis program at a final magnification of 80x (newborn) and 40x (100-d and 150-d old animals). Finally, the muscle fiber density and the total cross sectional area values of the semitendinosus muscle in the newborn animals were used to calculate the total muscle fiber number per sample. As the whole semitendinosus muscle in both 100-d and 150-d old is large and very heavy, it was unfeasible to manipulate them at the slaughter house. Therefore, the information on muscle area, and consequently total muscle fiber number, is missing in those experimental groups.

### Muscle fiber diameter

The diameter of the muscle fibers cross section [28] was determined using digital images randomly selected in the NB, 100-d and 150-d old pigs. Approximately 250 muscle fibers, chosen at random, from five animals of each experimental group, were measured resulting in a total of 1,250 fibers in the NW and IUGR groups. The diameter was measured using the analysis software Image J Software (1.49v – free version, National Institutes of Health, Bethesda, MD, USA).

### Volumetric density of muscular components

The volumetric density (%) of the muscular components, including muscle fibers, interstice, adipocytes and blood vessels for the three ages studied was obtained by point counting. Ten randomly selected sections per animal in each experimental group were examined under a light microscope (Olympus BX-51), with a 10x eyepiece fitted with a square lattice containing 284 intersections. Ten fields (total of 2,840 points) were randomly selected per animal at 400X magnification. The number of intersections on pertinent structures over the entire tissue section was counted by predetermined and systematic movement of sections across the grid without overlap.

Volume density of each muscular component was obtained by dividing the sum of points falling on each structure by the total number of points over the tissues. The results were expressed as a percentage of the muscle volume obtained by multiplying the volume density of each muscular component by 100.

### Immunofluorescence

To identify and quantify muscle fiber types (Myosin Heavy Chain - MyHC) type I and the isomeric forms type II (IIa, IIb and IIx) in the semitendinosus muscle, immunofluorescence was performed. Moreover, to evaluate the maturation of muscle fibers, the presence of the embryonic isoform (emb-MyHC) was also evaluated in NB, 100-d and 150-d old animals from both experimental groups. Myosin Heavy Chain is a sarcomeric protein, part of a complex which is essential for muscle development (embryonic isoform) and for slow (type-I) or fast (type-II) muscle contraction.

Briefly, sections were dewaxed, rehydrated and microwaved for 3 X 5 minutes in 0.1M sodium citrate buffer (pH 6.0) for epitope antigen retrieval, cooled down to room temperature and rinsed with phosphate buffered saline pH 7.4 (PBS). Blocking of reactive aldehyde groups was carried out with 100% (v/v) methanol for 30 minutes. Sections were maintained in Tween 20 solution (0.1% v/v in PBS) and blocked with 1% (w/v) of BSA in Dulbecco’s PBS for 30 minutes at 4°C. All samples were subjected to overnight incubation (12 to 15h at 4°C) with five different primary antibodies (**Table 1**). Negative controls were maintained without primary antibodies in PBS at 4°C. All sections were then incubated (90 min at 4°C) with their respective secondary antibody (**Table 1**). Finally, nuclei were stained using DAPI, and slides were mounted with glycerol 50% (v/v in Dulbecco’s PBS). For evaluation of slow and fast contraction muscle fibers, samples of rabbit semitendinosus muscle were used as positive and negative controls, according to the manufacturer, and for muscle fibers maturation, newborn pig tissues were used as positive and negative controls.

**Table 1.**
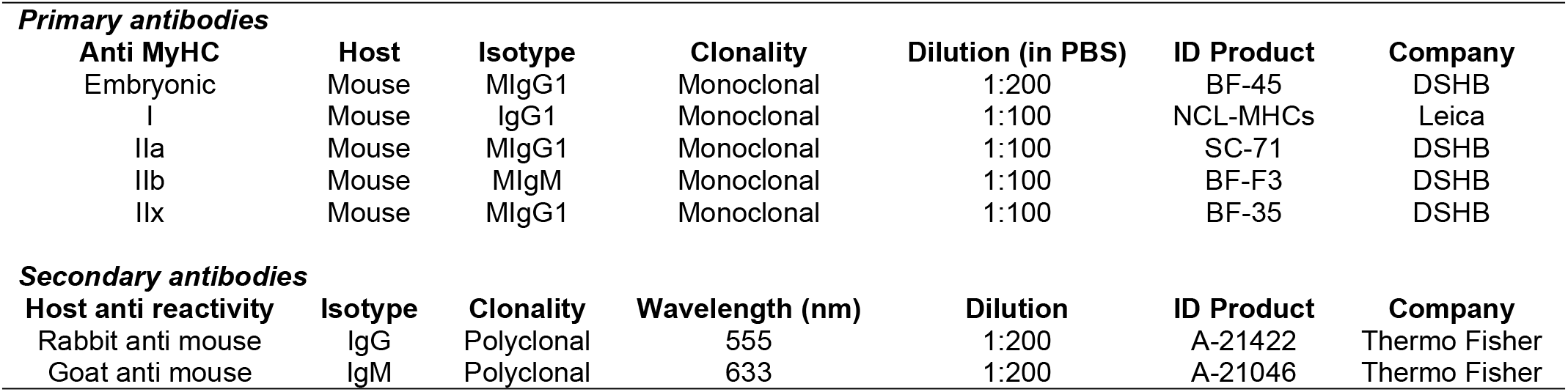
Description of primary and secondary antibodies used in the immunofluorescence analysis to identify immature (embryonic), slow (I) and fast (IIa, IIb and IIx) Myosin Heavy Chain (MyHC) protein isoforms in the pig semitendinosus muscle. IgG and IgM isotype primary antibodies were recognized by IgG and IgM isotype secondary antibodies, respectively.

The quantification of MyHC muscle fibers was obtained through the Zen software (Blue edition 2.3 – free version, Carl Zeiss Microscopy GmbH, 2011). The proportion of MyHC muscle fiber types was calculated by the ratio between the number of MyHC (-I, -IIa, -IIb) positive and the total number of muscle fibers (positive plus negative), in at least three fields of the semitendinosus muscle cross sections per animal. Since MyHC IIx fibers are not recognized by the antibody, the identification of these myofiber types was made considering only the negative cells, and further proportion was calculated as previously described. In the same fields, the evaluation of the MyHC fiber types’ diameter (Feret’s diameter - Dubach-Powell, 2011) was performed through the Image J software. Hybrid fibers were ignored.

### Relative expression of myogenic, adipogenic and growth-related genes by RT-qPCR

To understand the molecular mechanisms of postnatal skeletal muscle development, the relative expression of some myogenic, adipogenic and growth-related genes was quantified by RT-qPCR. The protein interaction network of those genes was established, using STRING database (version 10.5 – http://string-dg.org/) [29] with minimum required interaction score defined as higher confidence (0.700) (**Fig 1**).

**Fig 1.**
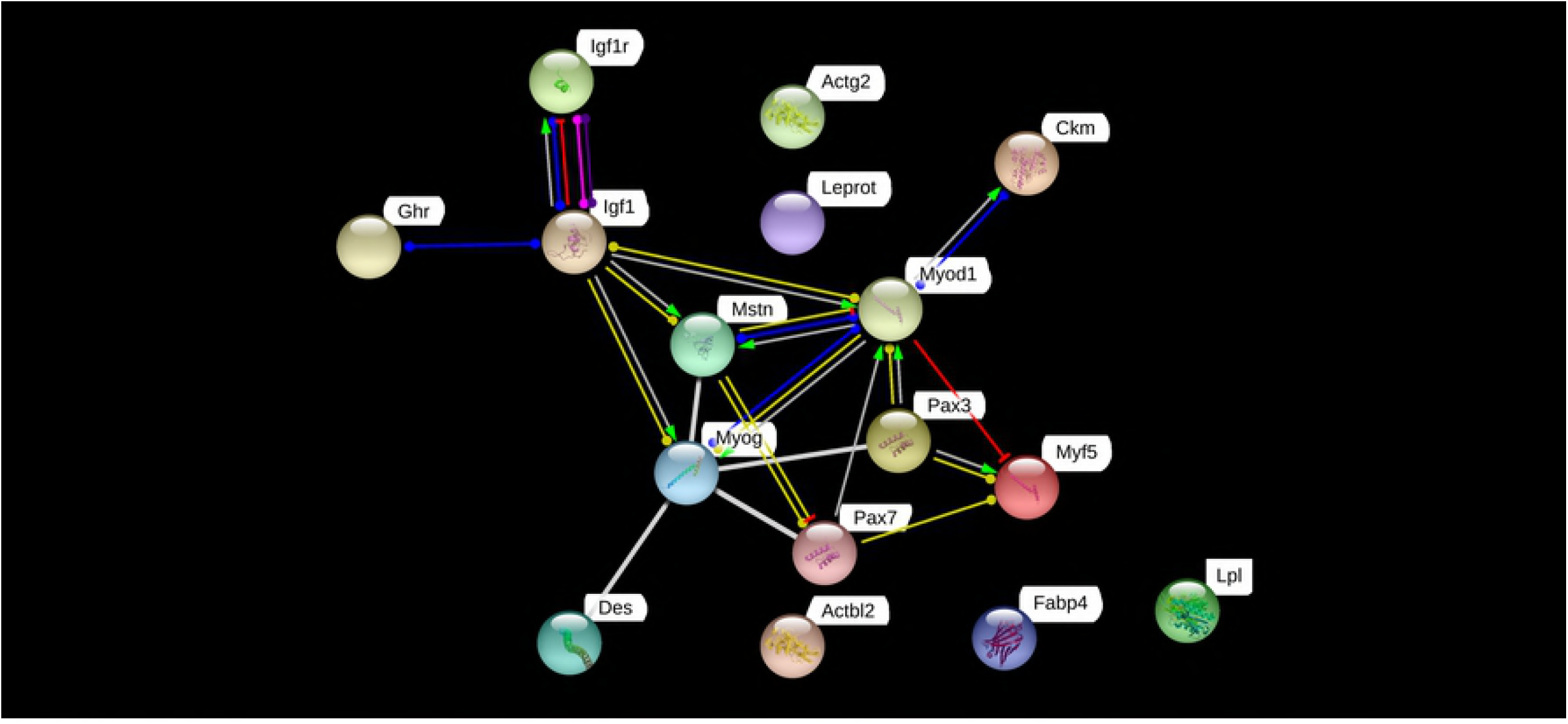
Predicted protein interaction network of the respective myogenic, adipogenic and growth-related genes involved in postnatal skeletal muscle development of higher mammalian. Nodes represent the proteins originated from their respective genes. When connected by edges, these proteins have known interactions. When isolated, these proteins have no physical or functional interactions.

Total RNA was isolated using Trizol^®^ Reagent (Life Technologies) following manufacturer’s instructions and its quality was evaluated using the Agilent RNA Screen Tape System and Agilent RGK Screen Tape System (Agilent Technologies, Mississauga, Ontario, CA) protocols. From each sample, 1 μg of total RNA was converted in cDNA, following the High Capacity cDNA Reverse Transcription Kit (Applied Biosystems – Life Technologies, Burlington, Ontario), from both oligo d(T) and random primers. Primers were designed based on pig mRNA sequences available at the Ensembl database (**Table 2**).

**Table 2.**
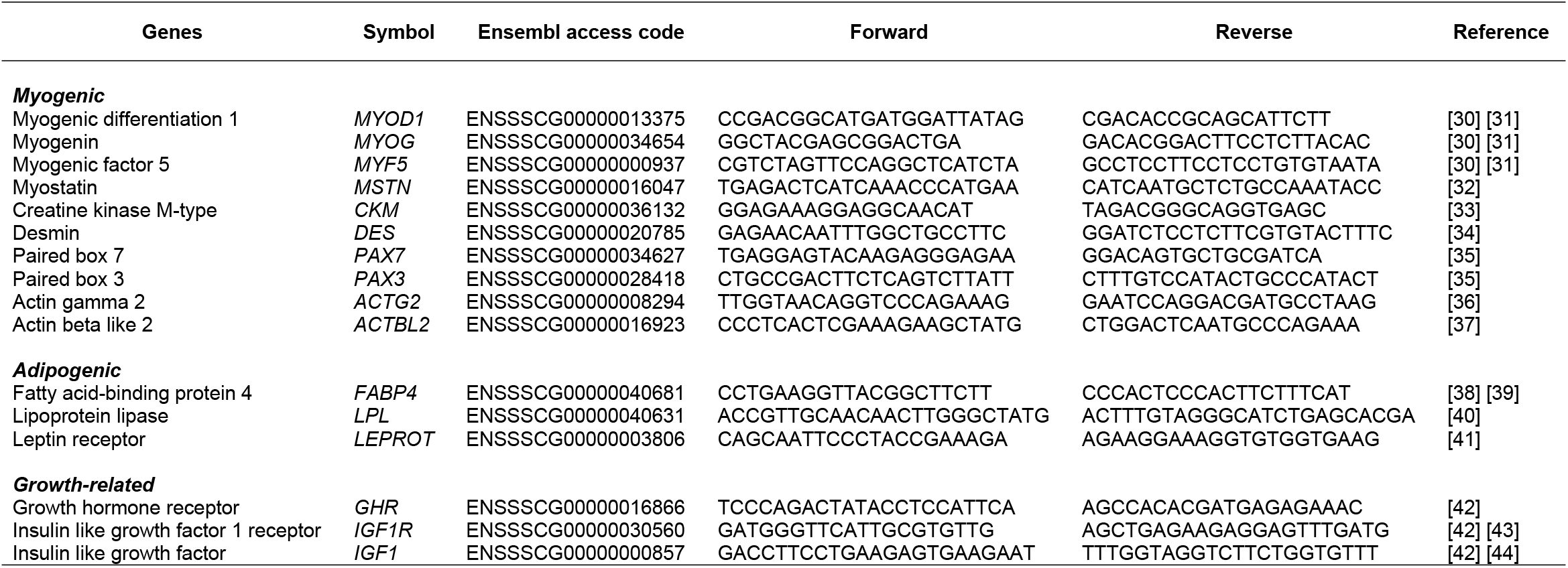
Myogenic, adipogenic and growth-related genes selected for evaluation by RTq-PCR, with their symbols, ensembl access code, forward and reverse sequence and reference.

RT-qPCR was conducted using Fast SYBR^®^ Green Master Mix (Applied Biosystems), 0.4-0.8 mM of each primer, and 1 μL of each 1:10 diluted cDNA, in a final volume of 10 μL. Reactions were performed in a 7900 Real-Time PCR Machine (Applied Biosystems), using the following cycling parameters: 50°C × 2 min, 95°C × 2 min, followed by 40 cycles of 95°C × 15 sec, 60-62°C × 30 sec, and 72°C × 20 sec. The dissociation step was performed at the end of the amplification step to allow the identification of the specific melting temperature for each primer set. All reactions were performed with technical duplicate and experimental triplicate (NW) or quintuplicate (IUGR). The calibration curve data such as the mean slope, intercept, PCR efficiency and R^2^ values are shown in **S1 Table**.

The relative gene expression analysis was calculated using the 2^-ΔΔCt^ method [45], and was analyzed from newborn (NB) until adulthood comparing, in each group, the three postnatal periods as follows: (i) postnatal myogenesis, where NB and 100-d samples were used; (ii) postnatal development, where NB and 150-d old samples were used; and (iii) hypertrophy, where 100-d and 150-d old samples were used. Results are expressed as the difference of the relative gene expression between ages.

### Statistical analysis

All variables measured were tested for normality prior to analyses, using the univariate procedure of the Statistical Analysis System (SAS Institute Cary, NC, USA, version 8.2). Data were analyzed as a randomized complete block design, each block consisting of two littermates. The statistical model included birth weight class and block as fixed factors and pig as random factor.

Treatment effects on growth performance, histomorphometrical analysis, and muscle fiber types were analyzed using the general linear model (GLM) procedure of SAS. Least square means were compared using the Student’s t-test with P < 0.05 being considered significant. In the tables and figures, data are reported as least square means and the pooled SEM. The relative expression of myogenic, adipogenic and growth-related genes were analyzed using the Relative Expression Software Tool (REST 2009, QIAGEN).

## Results

### Newborn group data and postnatal growth performance

The organs weights and the brain to liver weight ratio of the newborn piglets are shown in **Table 3**. All organs of NW piglets, including the brain, were heavier than the organs of their IUGR counterparts (P < 0.01). The brain to liver weight ratio was increased in IUGR piglets (P < 0.05), which confirms the occurrence of intrauterine growth restriction. NW animals presented higher body weight at birth (**Fig 2**), and these differences were maintained throughout the postnatal period (weaning, nursery and grower/finisher) (P < 0.05).

**Table 3.** Organ weights (g) and brain/liver weight ratio in normal weight (NW) and intrauterine growth restriction (IUGR) newborn piglets.

| Parameters | Experimental groups |  | SEM |
| --- | --- | --- | --- |
|  | NW | IUGR |  |
| N | 10 | 10 |  |
| Birth weight, kg | 1.5<sup>a</sup> | 0.9<sup>b</sup> | 0.01 |
| Small intestine | 62.6<sup>a</sup> | 36.6<sup>b</sup> | 2.10 |
| Large intestine | 16.8<sup>a</sup> | 10.6<sup>b</sup> | 0.30 |
| Liver | 43.8<sup>a</sup> | 23.9<sup>b</sup> | 1.20 |
| Heart | 10.5<sup>a</sup> | 6.7<sup>b</sup> | 0.25 |
| Brain | 28.7<sup>a</sup> | 25.2<sup>b</sup> | 0.40 |
| Stomach | 9.0<sup>a</sup> | 5.8<sup>b</sup> | 0.30 |
| Brain: Liver | 0.7<sup>a</sup> | 1.1<sup>b</sup> | 0.04 |

<sup>a,b</sup> Within a row, LSmeans with different superscripts differ ( $P < 0.05$ ).

**Fig 2.**
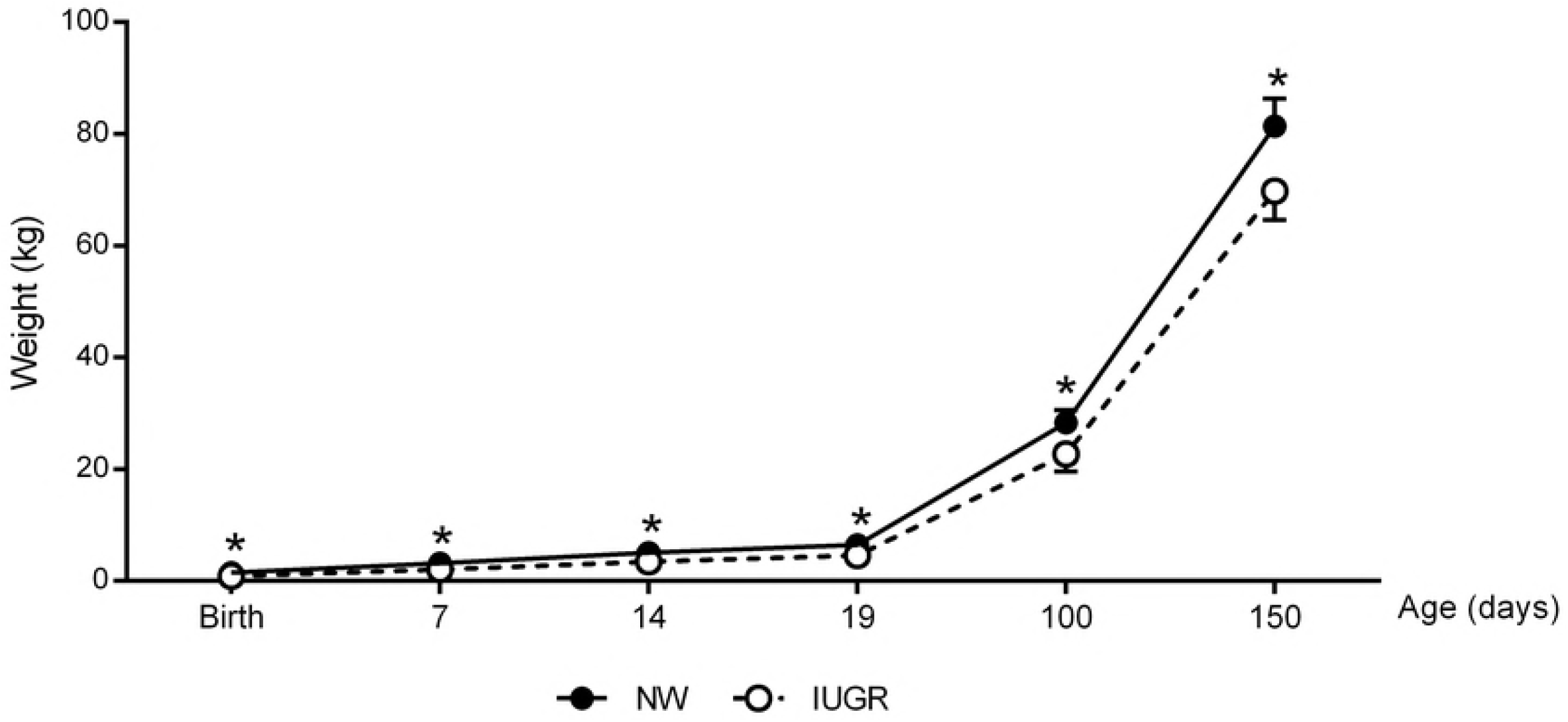
Postnatal growth curve (kg) from normal weight (NW) and intrauterine growth restricted (IUGR) gilts. * P < 0.05

### Histomorphometrical analysis

The histomorphometrical analyses of the semitendinosus muscle in both experimental groups at the ages evaluated are summarized in **Table 4**. These analyses revealed that growth restriction *in utero* did not affect the proportions of muscular components in the NB group (P > 0.05). On the other hand, NW newborn pigs had greater muscle fiber diameter and consequently lower fiber density (fibers/mm^2^), compared to their IUGR littermates (P < 0.01). In addition, the cross-sectional area of the semitendinosus muscle was smaller in IUGR females from the NB subgroup (P < 0.05).

**Table 4.**
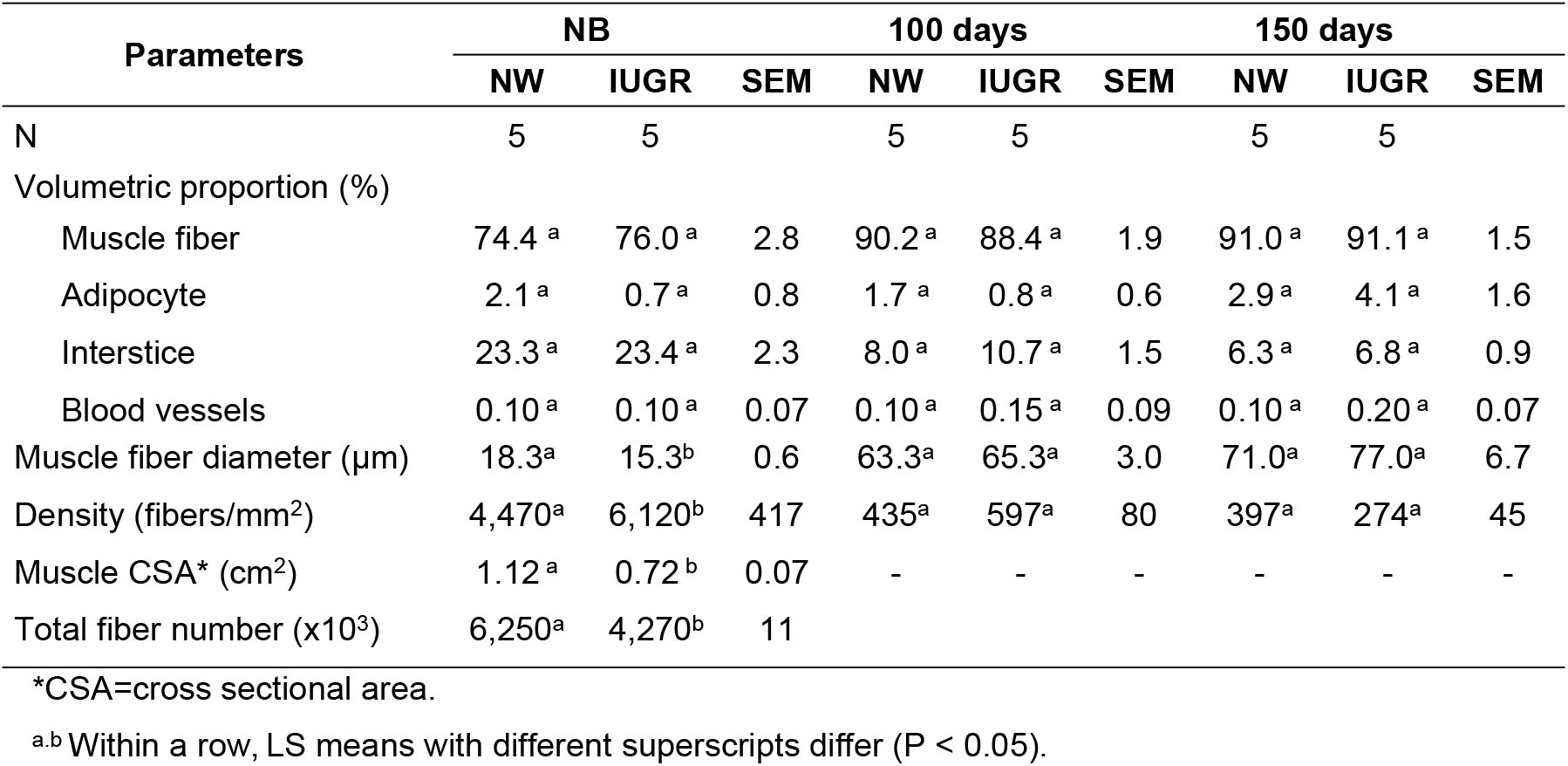
Histomorphometrical analyses of the semitendinosus muscle of newborn, 100-d and 150-d old pigs from normal weight (NW) and intrauterine growth restricted (IUGR) experimental groups.

Even though muscle fiber diameter and density as well as muscle cross-sectional area were different in the newborn animals, these differences were not apparent over time, as both NW and IUGR animals showed similar muscle fibers diameter and density at 100 and 150 days of age (P > 0.05).

### Evaluation of muscle fiber types

The NW group presented emb-MyHC protein isoform only at birth; this fiber type was absent at the later stages of postnatal development (**Table 5; Fig 3**). However, IUGR animals presented different fiber type composition, as emb-MyHC could also be detected in 100-d and 150-d old females (**Figs 3B and D**). At birth, the IUGR piglets showed almost the double of the proportion of MyHC-I (slow-twitch oxidative) fibers compared to their NW counterparts, and yet this difference has disappeared over time, as this fiber type frequency was similar in both 100-d or 150-d old gilts (**Figs 3E-F** and **3G-H**). Regarding MyHC-II (fast-twitch glycolytic) muscle fibers, neither types -IIa, -IIb or -IIx were affected by intrauterine growth restriction at 100-d of age. Interestingly, the proportions of MyHC-IIa and -IIb were numerically higher in IUGR females at 150-d of age but did not reach the level of significance (p=0.059 and p= 0.087, respectively). However, growth restriction *in utero* affected the proportion of MyHCII-x, which was lower in IUGR gilts (**Figs 3Q-R** and **3S-T**). Furthermore, the hypertrophy capacity evaluated between 100-d and 150-d old pigs revealed that it was lower in IUGR females in all MyHC fiber types evaluated (**Table 6**).

**Table 5.**
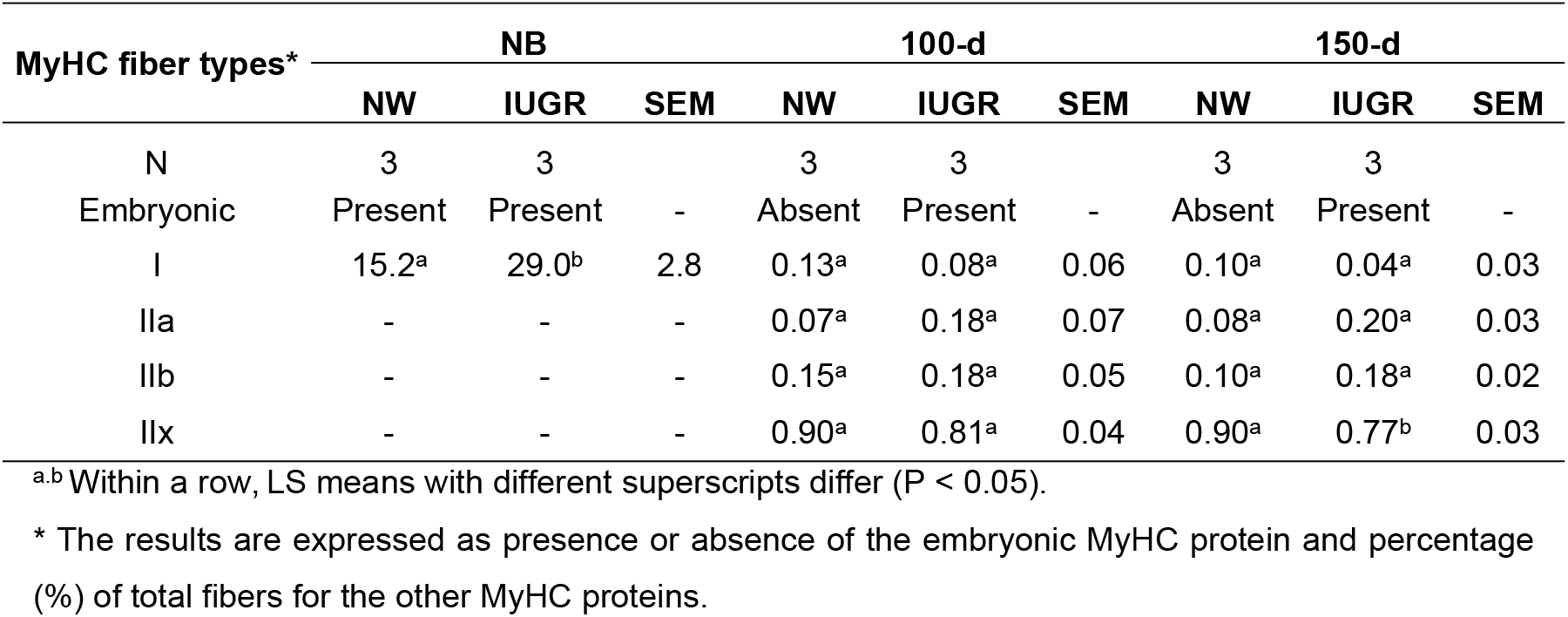
Quantification of Myosin Heavy Chain (MyHC) fiber types in the semitendinosus muscle in newborn (NB), 100-d and 150-d old gilts from normal weight (NW) or intrauterine growth restricted (IUGR) experimental groups.

**Table 6.** Hypertrophy based on the Myosin Heavy Chain (MyHC) fiber types in the semitendinosus muscle of 100-d until 150-d old pigs from normal weight (NW) and intrauterine growth restricted (IUGR) animals.

| MyHC | NW |  |  | IUGR |  |  | Hypertrophic capacity of IUGR related to NW (%)* |
| --- | --- | --- | --- | --- | --- | --- | --- |
|  | 150-d | 100-d | Hypertrophy (150d - 100d) | 150-d | 100-d | Hypertrophy (150d - 100d) |  |
| I | 3.47 | 1.12 | 2.35 | 1.84 | 1.33 | 0.51 | 22 |
| IIa | 3.24 | 1.47 | 1.77 | 2.98 | 1.37 | 1.60 | 91 |
| IIb | 3.34 | 1.46 | 1.88 | 2.55 | 1.61 | 0.94 | 50 |
| IIx | 3.87 | 1.65 | 2.22 | 3.33 | 1.99 | 1.33 | 60 |

\*The hypertrophic capacity of IUGR pigs is based on the same parameter of their NW counterparts.

**Fig 3.**
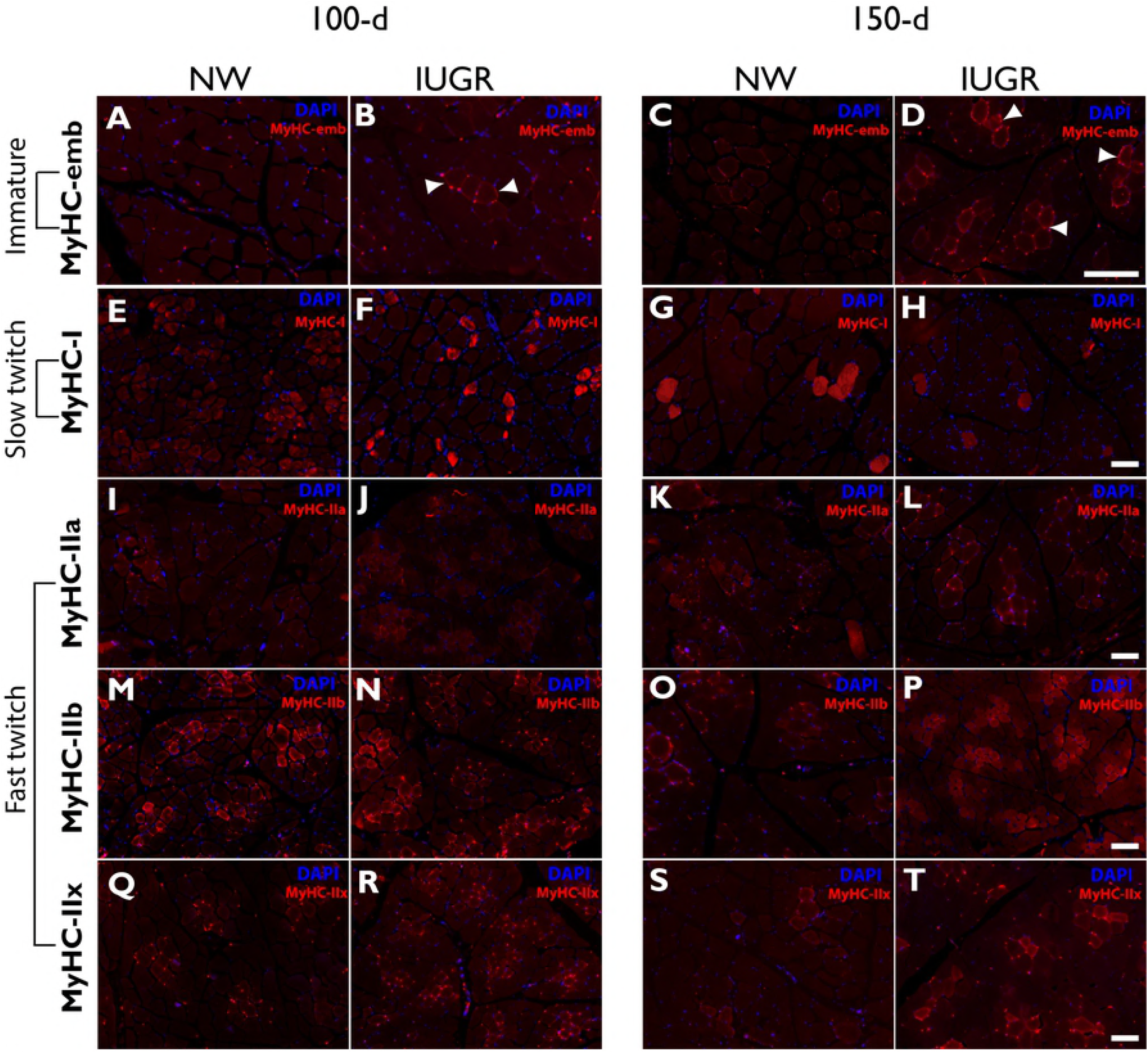
Immunostaining of immature (embryonic – emb), slow (I) and fast (IIa, IIb and IIx) twitch Myosin Heavy Chain (MyHC) protein isoform in the semitendinosus muscle from normal weight (NW) and intrauterine growth restricted (IUGR) pigs at 100-d and 150-d old. The bars represent 96 μm.

### Relative expression of myogenic, adipogenic and growth-related genes in the semitendinosus muscle during postnatal development

The relative gene expression data revealed that in NW animals only *IGF1* was found to be up-regulated (P < 0.001) in the skeletal muscle from 100 d-old animals, compared to the NB ones (postnatal myogenesis; **Fig 4A**). No other significant differences were observed in the expression of the genes associated with muscle development and homeostasis (*PAX3, PAX7, MYOD1, MYF5, MYOG, MSTN and CKM, DES, ACTG2, ACTBL2*), growth (*GHR* and *IGF1R*) and lipid metabolism (*FABP4, LPL* and *LEPROT*) between NB and 100 d-old samples. During the same period, however, IUGR skeletal muscle samples presented a different expression pattern: *MYOD, MYOG, MYF5, DES* and *PAX7* were all found to be down-regulated in 100 d-old samples, compared to the NB (**Fig 4A’**); while no significant differences were found in the other gene expression ratios.

**Fig 4.**
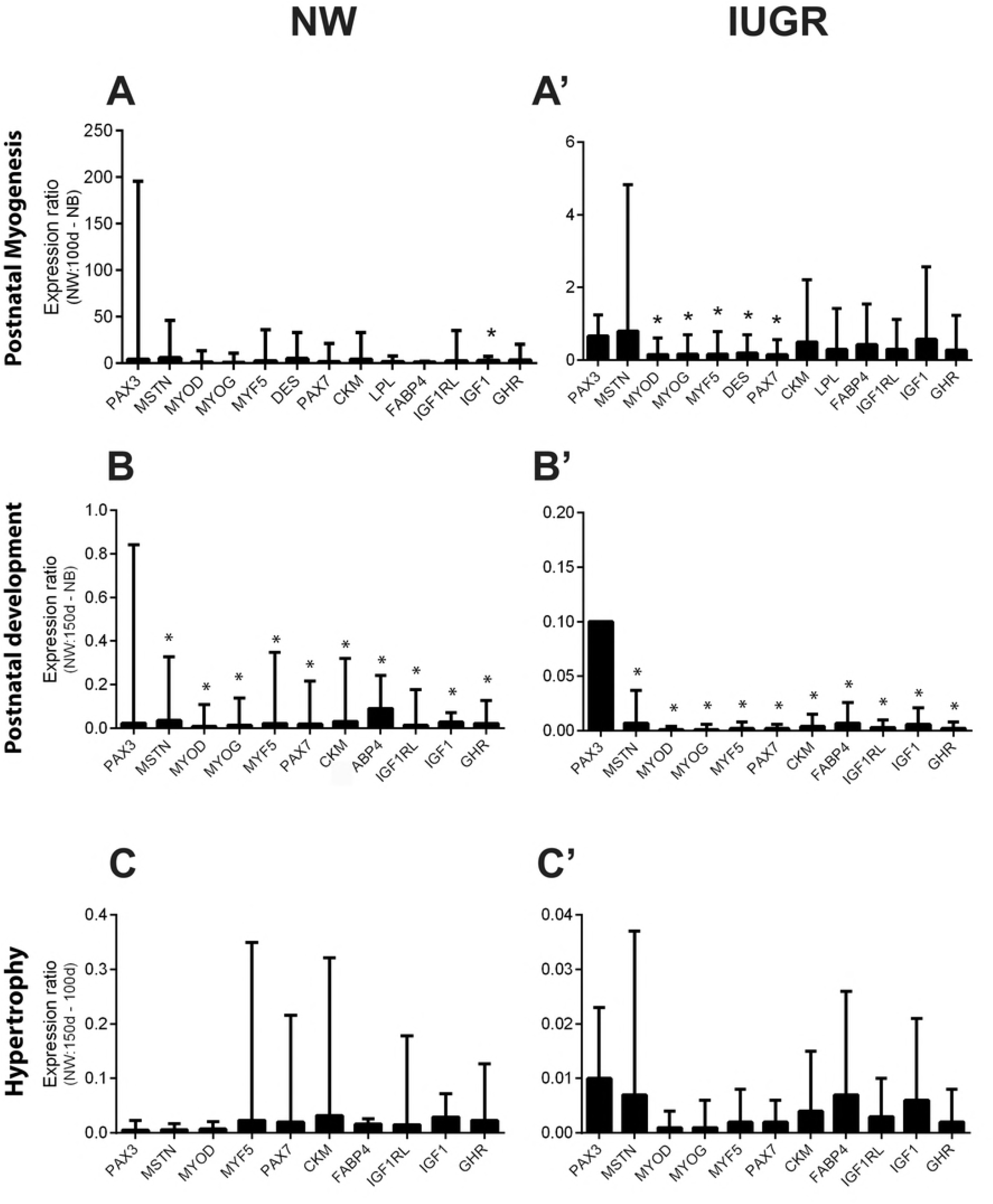
Gene expression evaluation from normal weight (NW) and intrauterine growth restriction (IUGR) pigs in three postnatal life phases: myogenesis (A-A’), postnatal development (B-B’), and hypertrophy (C-C’).

The analysis between 150 d-old and NB skeletal muscles (postnatal development) revealed a similar expression pattern for both NW and IUGR: all genes were found to be down-regulated in both samples, excluding *PAX3* in NW animals, which showed similar expression between those ages (**Fig 4B-B’**). However, the repression levels observed in IUGR samples were always at least five times higher than the ones observed in NW (i.e. *MSTN* expression level in the NW animals was 0.036 - P<0.001; while in IUGR was 0.007 - P<0.003). The same tendency of gene repression was observed when the comparison was performed between 100 and 150 d-old samples (hypertrophy; **Fig 4C-C’**).

## Discussion

Several studies have investigated skeletal muscle alterations due to intrauterine growth restriction, and yet there is little information on postnatal myogenic program and its structural changes, including fiber type, to further elucidate the effects on postnatal muscle growth. It is well established that total muscle fiber number, which is determined during fetal stage, is a crucial component of postnatal growth, and postnatal skeletal muscle development is mainly due to the increase in muscle fiber size, with no net formation of new muscle fibers [46]. Since the number of muscle fibers formed during fetal stage is dependent on the number of available myogenic progenitor cells and their proliferation is highly sensitive to nutrient availability in utero [46], a better understanding of the impact of intrauterine growth restriction on skeletal muscle development is important.

Hence, the present study investigated morphofunctional alterations due to intrauterine growth restriction on skeletal muscle postnatal development in gilts and their possible postnatal myogenic program origins. In particular, it was shown that IUGR promotes structural changes in muscle fibers and alters the proportion of MyHC-I myofibers at birth, with smaller muscle fibers diameters and increased proportion of MyHC-I myofibers in IUGR animals, whose phenotype is related to postnatal myogenic program impairment. However, these structural changes did not persist over time and this may reflect a particular characteristic of the myofiber which is plasticity. To the best of our knowledge, this is the first report showing the skeletal muscle morphological phenotype and the gene expression profile of skeletal muscle development from birth to adulthood in IUGR individuals, using the pig as an experimental model.

Similar to previous studies [17] [47] where growth rates of different birth weight pigs were investigated, IUGR gilts showed lower body weights throughout the postnatal development period. This may be due to the lower muscle fiber number observed in IUGR piglets at birth, as a positive correlation between muscle fiber number and performance traits has been previously reported [48] [49]. Besides the low number of muscle fibers, the smaller muscle fiber diameter and muscle cross sectional area observed in the IUGR group at birth may also contribute to the low growth rates reported in the IUGR females during all ages evaluated. Despite the differences in the semitendinosus histomorphometry at birth, the same parameters were not apparent at 100-d and 150 days of age, suggesting that muscle hypertrophy is not affected by IUGR. Indeed, similar pattern in the relative expression of genes during the hypertrophy period (100 to 150 days of age) confirms this finding.

Regarding the quantification of muscle fiber type, the proportion of MyHC-I myofibers was apparently not compromised by growth restriction *in utero* at 100-d and 150-d of age. These findings are in agreement with Choi and colleagues [50], which did not report any differences in the proportion and area of MyHC-I fibers relative to live weight at slaughter. Since higher proportion of MyHC-I fiber types contribute to superior meat quality [51], the present results suggest that IUGR may not have direct consequences for meat sensory quality characteristics. Indeed, the impact of birth weight on meat quality has been recently investigated by Alvarenga and colleagues [52], who reported a higher shear force in the Longissimus dorsi muscle from IUGR barrows. Interestingly, the presence of the embryonic MyHC isoform in the semitendinosus muscle of 100-d and 150-d old IUGR animals suggests muscle immaturity, once it was absent in NW animals at the same age. Hence, the presence of embryonic MyHC may impair muscle functionality in IUGR pigs. Our results corroborate with those reported by Perruchot and colleagues [20] which demonstrated greater amount of the embryonic MyHC isoform in small fetuses, suggesting delayed myofiber development.

The molecular regulation of skeletal muscle postnatal myogenesis in IUGR animals still lack clarification, especially regarding postnatal development. In this regard, at the same age, while skeletal muscles from NW animals showed the up-regulation of *IGF1* expression, IUGR ones showed the down-regulation of the classic *PAX7, MYF5, MYOD* and *MYOG* myogenic regulatory factors (MRFs) and the cytoskeleton component *DES*. These results revealed the strong difference in the expression pattern of myogenic markers between these samples with NW animals inducing the expression of an important growth factor for skeletal muscle cells and the IUGR delaying the expression of signals that induces skeletal muscle myogenesis. Since the cells responsible for the expression of these markers were originated during the prenatal period, they may have been influenced by the uterine environment, as the embryo-fetal stage is crucial for skeletal muscle development [46]. In this sense, insults during prenatal development may affect gene expression related to myogenesis through DNA methylation which will have long lasting effects on postnatal development [26].

Since *PAX7* participates in regulating the behavior of skeletal stem cells [53], its deficiency might play an important role in myogenesis impairment. During postnatal development, *PAX7* is under expressed in IUGR animals, but not absent. Hence, it is reasonable to interpret that the recruitment of satellite cells to the myogenic lineage by *PAX7* was less intense. Consequently, *MYF5* and *MYOD* were under expressed and *MYOG*, a marker of terminal differentiation of the muscular lineage [54] as well as *DES*, were also under expressed. Therefore, it is suggested that the low expression of *PAX7* affected the myogenic program causing the early closure of postnatal skeletal muscle myogenesis. This was shown by the immune fluorescence determination of the embryonic MyHC protein in the skeletal muscle tissue in 100-d and 150-d old IUGR pigs.

Finally, the presence of the embryonic MyHC protein isoform in IUGR adult muscle associated with the description of the myogenic regulatory factors deserve attention. The myogenic regulators described herein become candidates in the investigation of postnatal myogenic program to ensure the proper development of skeletal muscle in swine breeding programs and in human health. Although the conditions imposed by the intrauterine environment may cause skeletal muscle morphofunctional changes in IUGR individuals, future studies are necessary to investigate the mechanisms which limit postnatal muscle growth and develop strategies that might prevent the onset of diseases in later life.

## Acknowledgements

The authors gratefully acknowledge Professor Cristina Guatimosim Fonseca from the Department of Morphology – ICB/UFMG for donating the antibodies and her scientific support during the immunofluorescence analyses and Joan Turchinsky (AFNS; University of Alberta) for the technical support she provided as a lab technologist for the research group.

## Supporting information

**S1 Table. qPCR calibration curve data.** The slope, intercept, efficiency and R2 are presented.

